# Distinct neocortical and entorhinal networks for time, space, and reward

**DOI:** 10.1101/2025.10.13.682150

**Authors:** John B. Issa, Daniel A. Dombeck

## Abstract

Variables relevant to behavior and cognition are often synthesized via computations that combine external cues and internal states. Especially salient and ubiquitous among these variables are time, space, and reward, but how they are represented throughout the neocortex and the entorhinal cortex is not well understood. Here, we utilized a behavioral task in virtual reality that includes timing, navigation, and reward components. Using a multifocus microscope that permits simultaneous mesoscopic Ca^2+^ imaging of neocortex and entorhinal cortex, we uncover dedicated and distinct subnetworks for interval timing, spatial navigation, and reward. Interestingly, the timing subnetwork demonstrated more prominent signaling in expert mice and features a strong contribution from barrel cortex that is not explained by ongoing whisking, pointing towards a role for this cortex in interval timing. Our results provide previously unknown details about the brainwide networks related to time, space, and reward and will allow for future interrogation of their interactions and computations.

## Introduction

The neocortex handles a diversity of information streams^1^, with unimodal sources of information such as vision, touch, and movement represented in primary sensory and motor regions of neocortex. However, variables relevant to behavior and cognition are often not explicitly represented in these unimodal signals and must instead be synthesized within the dynamical systems of the brain via computations that combine external cues and internal states^2–4^. Variables such as time, reward state, and spatial position may be especially relevant to decision-making and memory^5^. They are known to be represented in the hippocampus, which contains time cells^6^, reward cells^7^, and place cells^8^, which serve as building blocks for episodic memories^9^. Where do these signals arise from, and how does the relevant information reach the hippocampus? The primary inputs to the hippocampus arise from the entorhinal cortex, which itself is extensively connected to much of the neocortex^10^. How are correlates of timing, reward, and spatial navigation distributed across the neocortex and the entorhinal cortex^11^?

Here, we built a custom multifocus microscope, which permits mesoscale Ca^2+^ imaging of much of the neocortex along with the entorhinal cortex in the same preparation. We used a behavioral task in virtual reality that serves to isolate the aforementioned variables of time, reward state, and spatial position. We wished to determine which regions in neocortex and entorhinal cortex are involved in representing these variables. Using nonnegative factorization and regression methods, we isolate signals related to each individual variable, uncovering distributed subnetworks that span large regions of neocortex and entorhinal cortex. We also find that, as mice gain expertise on the timing component of the task, the timing subnetwork grows stronger, consistent with the capability of cortex to generalize and learn arbitrary computations^12^.

## Results

### Simultaneous imaging of neocortex and entorhinal cortex during a multi-component task

We utilized a task that engages three different aspects of behavior: reward, spatial navigation, and interval timing^13^ (Figure 1A). Head-fixed mice ran on a linear treadmill to traverse a 3.1-m virtual track. A water reward was delivered at the end of the track, followed 4 seconds later by teleportation back to the start of the track. The timing portion of the task was achieved by placing a door halfway down the track where mice need to stop running for the target duration (typically 6 seconds), at which point the door opens and the mice can proceed down the track. However, if they resume running before the target time, the timer resets until the next attempt at running. Once mice learned the task, the door was made invisible so that no cues about the timing interval were explicitly provided. Early in training, wait times followed an exponential distribution, reflecting a Poisson process, but in well-trained mice wait times followed a normal distribution with a mean slightly below the target time, possibly reflecting information gathering and optimal foraging^14^. We classified mice as “novice” or “expert” based on the distribution of wait times being either exponential or Gaussian, respectively (Figure 1A). Importantly, the timing portion of the task requires the mouse to stand still on the treadmill for the target waiting time; thus, while some motor behaviors could still be ongoing, large movements and outright running are eliminated, helping to isolate timing-related signals that would otherwise be overwhelmed by velocity-related signals in the neocortex^15^.

**Figure 1.**
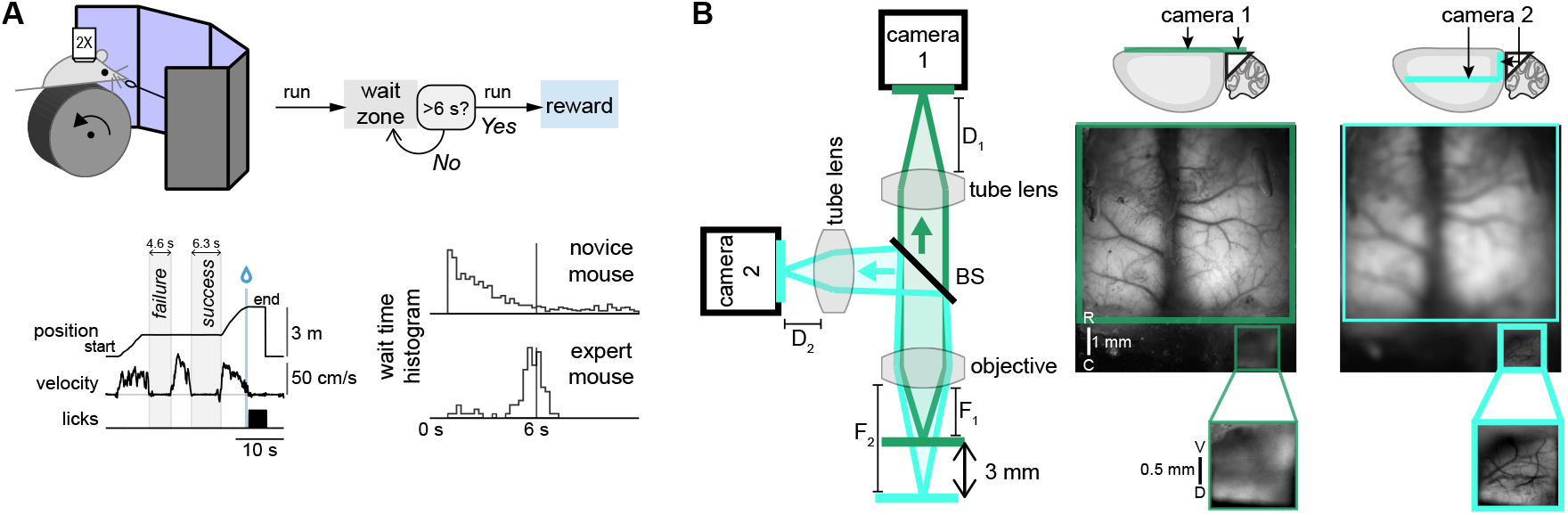
Behavior and imaging setup. A) Mice learn a VR task with three components: 1) run on a treadmill to travel down a 3.1 m virtual track; 2) receive reward at the end of the track (indicated by vertical blue line); 3) halfway down the track, stop running at an invisible door for at least 6 s before they can continue running to the end of the track. In example shown, first attempted stop is a failure since it is less than 6 s, but the second attempt is successful. In novice mice, histogram of wait times follows an exponential distribution while in expert mice it is normally distributed with a mean near the target time (0.25-s bins, with waits less than 1 s excluded). Multifocus microscope for multiplane imaging. Two imaging collection paths (formed by 50:50 beamsplitter, BS) go to two cameras, each with an adjustable distance to its tube lens, allowing independent control of the focal plane of each camera. Camera 1 is focused on dorsal cortex (green path) while camera 2 is focused on the entorhinal cortex (cyan path), as imaged through a microprism and typically ∼3 mm deeper than the first imaging plane.

Given that the entorhinal cortex may be involved in interval timing^13,16–18^, we wished to record signals from the entorhinal cortex and multiple cortical areas (including visual, somatosensory, and motor cortices) during this behavior. We used transcranial mesoscale Ca^2+^ imaging since this method can accommodate a large imaging field^19^. The low temporal resolution (on the order of hundreds of milliseconds) is acceptable for our task since task-relevant timescales are on the order of seconds. We used a transgenic GCaMP6s mouse with high expression levels throughout neocortex and entorhinal cortex^20^ (Supplementary Figure 1). To image the entorhinal cortex simultaneously, we developed a hybrid preparation that combines an implanted microprism (1.5 or 2 mm) over the entorhinal cortex^21^ along with a transcranial clear skull preparation for dorsal neocortex covered by an 8 mm coverslip. The imaging planes for these two windows are offset by ∼3 mm. To achieve simultaneous in-focus imaging of both regions, we built a multifocus fluorescence microscope (Figure 1B and Supplementary Figure 2). Using two imaging paths leading to two separate cameras, we manually adjusted the back focal length of the second camera, allowing us independent control of the focal planes of the two cameras. Two LEDs (470 nm and 405 nm) were alternatively pulsed (Supplementary Figure 3), allowing correction for hemodynamic changes^22^.

Typical imaging sessions lasted 20-30 minutes, during which trained mice successfully completed ∼30 laps. Individual imaging fields were registered to bregma and normalized to bregma-lambda distance. Neural activity was measured by computing ΔF/F_0_ and then deconvolving to recover an estimate of instantaneous firing rates (details for each processing step, including isosbestic correction and calculation of baseline fluorescence, are provided in the Methods). Fluorescence traces from seven exemplar 0.1 x 0.1 mm superpixels are shown for a 2-minute stretch of an imaging session (Figure 2A-B), along with task-relevant behaviors of treadmill velocity, licking, and position on the virtual track. Each of these signals is averaged across trials for three aspects of the task (Figure 2C): first, timing at the door, which is formed by normalizing time to the duration from the start to the end of the timing period; second, time to reward, which is aligned to the time of reward delivery and followed after 4 seconds by teleportation to the start of the track; third, track position, which spans the 3.1-m track distance and, at each 1-cm bin, averages across any times when the mouse is moving down the track. These trial-averaged signals provide a signature of the unique response properties of each pixel with respect to each aspect of the task. For example, pixel 1 ramps up during timing at the door and remains high during reward consumption, while pixel 2 is less active during timing but peaks during reward approach. Together, the multi-component behavior and multi-focus imaging yielded rich, high-dimensional readouts of motor activity, task performance, and cortical signaling.

**Figure 2.**
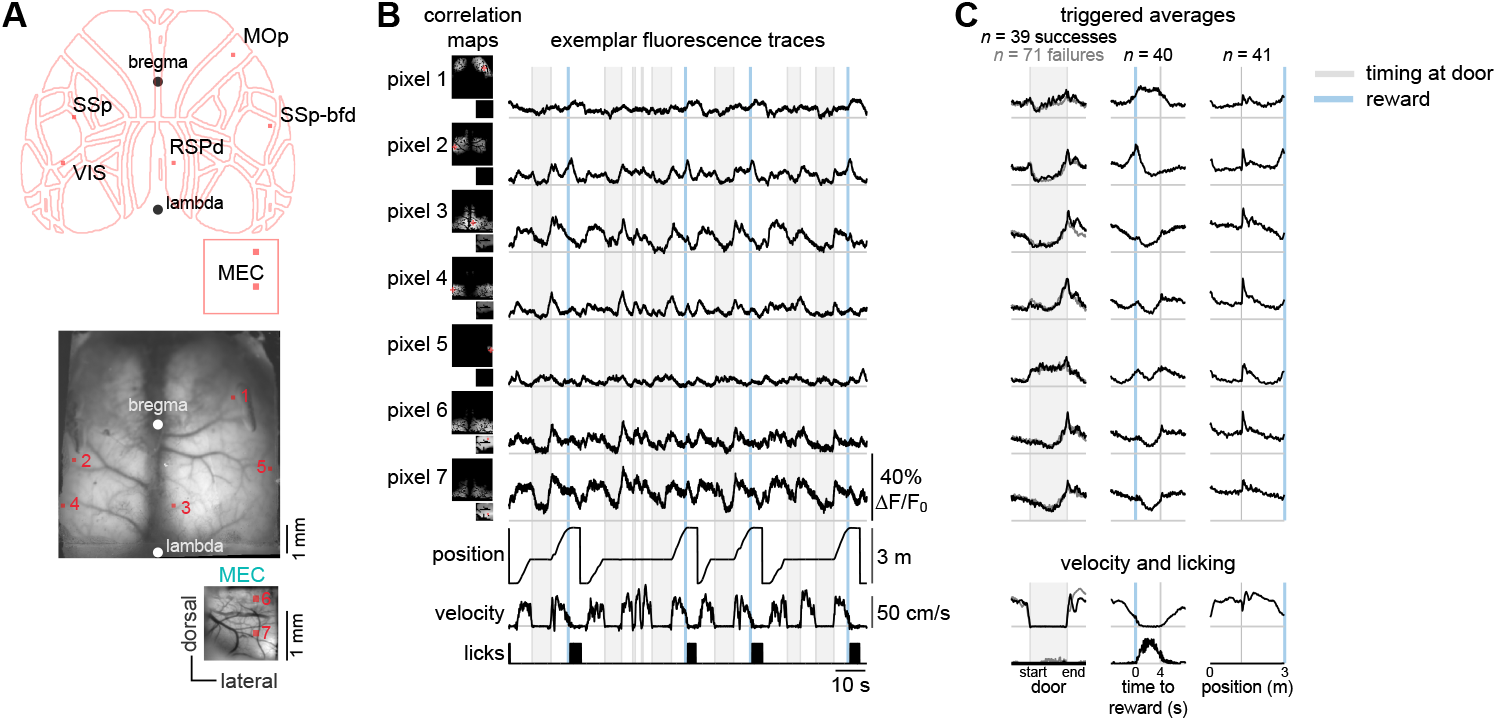
Brainwide Ca^2+^ signals during multi-component behavior. A) Mean fluorescence images acquired by two cameras, one focused on and cropped to the ∼8 mm window over dorsal neocortex and the other to medial entorhinal cortex (MEC). Exemplar pixels to be scrutinized in next panels are indicated by red dots. Movies are registered to bregma and lambda, enabling alignment to the 2D “flatmap” of the Allen Mouse Brain Common Coordinate Framework^43^. B) Fluorescence signals shown for a 2-minute period for the seven exemplar pixels, with behavior underneath. Thumbnails show correlation maps between each pixel and the remaining image. C) Triggered averages for the seven exemplar pixels and for velocity and licking with respect to the timing period at the door, the time to reward, and track position. For the timing period, time was normalized to the start and end of each wait attempt; further, trials were split into successes (wait time greater than the target time, which here was 6 s; dark trace) and failures (wait time greater than 2 s but less than the target time; gray trace). For time to reward, the teleportation back to the start of the track happens at 4 s, as indicated by the thin vertical gray line. For track position, only times when the velocity is positive are used; signals are resampled to 1-cm bins.

### Identification of task-relevant subnetworks

To explore differences across brain regions more systematically, we sought a method that would allow us to combine results across mice and, through dimensionality reduction, uncover a small number of components that are sufficient to explain a large amount of the variance in the fluorescence signals. With this goal in mind, we developed a group-wise non-negative matrix factorization algorithm^23^ (g-NNMF, Figure 3A). For each pixel from each imaging session, we formed a single vector by concatenating the trial-averaged signals for timing, reward, and track position. This vector has a length of 794 data points. Then we formed a matrix for all 36318 pixels across 9 imaging sessions, where each row is the trial-averaged vector for that pixel, resulting in a 36318 x 794 matrix **X**.

**Figure 3.**
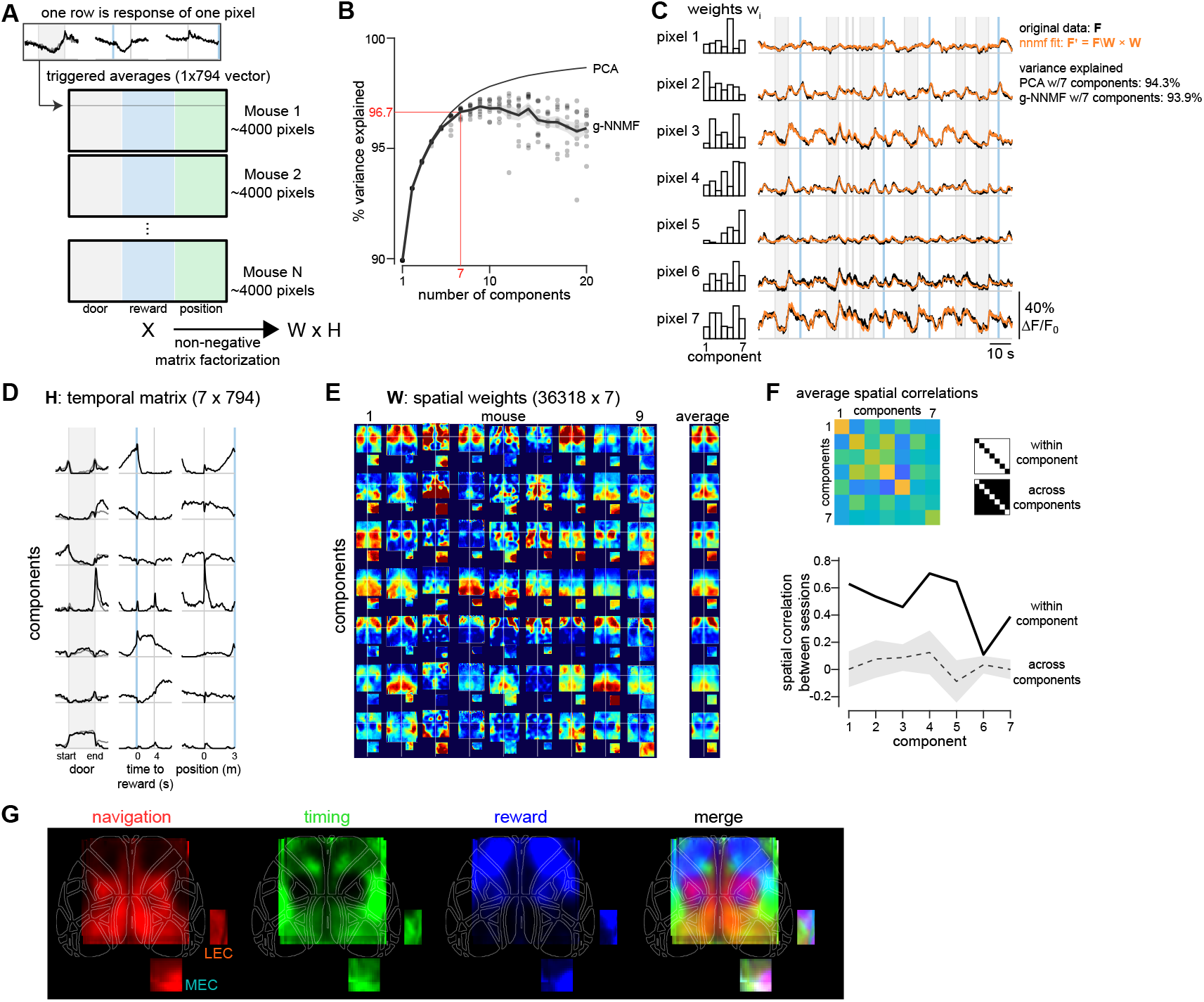
Isolation of task-relevant components using group non-negative matrix factorization. A) A multi-session matrix X is formed for all pixels across nine sessions (each from a different mouse). Each row corresponds to the triggered average of a single pixel, formed by concatenating triggered averages for timing, reward, and track position as shown in Figure 2C. B) Variance explained by non-negative factorization as a function of the number of components. Dots represent the 10 repeats for a given number of components; solid line and gray represent mean and standard error. For comparison, variance explained as a function of number of components is shown for principal component analysis (PCA). C) Weights (left) and reconstruction (right, in orange) using non-negative factorization with seven components is shown for the exemplar traces from Figure 2B. D) Temporal matrix H for the non-negative factorization of matrix X into H and W. E) Spatial weights W for the non-negative factorization. These are reshaped as 2D spatial maps for individual sessions, which are aligned to bregma. Mean weights (averaged across sessions) are shown as well. F) Spatial correlation between pairs of spatial weights for individual components and sessions, averaged either within component (same component, different sessions) or across components (different components, different sessions). G) Mean spatial maps across multiple components. Navigation: components 2, 3, and 6; timing: component 7; reward: components 1 and 5. Averages are across nine sessions (eight for MEC, one for LEC).

We chose the number of components to be seven, based on the elbow in variance explained as we increased the number of components (Figure 3B). Applying g-NNMF to **X**, we uncovered seven components, each of which has one row in the temporal matrix **H** (length 794) and one column in the weight matrix **W** (length 36318), the latter of which can be used to form 2D spatial maps for individual sessions (Figure 3C-E). Applying W to an individual session’s full movie **F** (which has 43432 time points and 3968 pixels) allows us to compress the movie to seven dimensions (43432 x 7), where each dimension corresponds to one component. Then, using the same weights, we can calculate an estimate of the full movie, **F’**. This recovered movie produces a good fit of the original movie, explaining 94% of the variance in the traces, nearly identical to factorization using principal component analysis (PCA), thus indicating that g-NNMF is optimally capturing variance with a low number of components (Figure 3C). We validated the factorization by holding out one session at a time, applying PCA or g-NNMF to the remaining sessions, and then testing the variance explained of the held out session. Both PCA and g-NNMF performed equally well (95.6% for PCA, 95.5% for g-NNMF; *p* = 0.07 with Wilcoxon signed rank test). Note that this factorization does not use any information about where each pixel came from, nor the temporal order of the individual points within the trail-averaged vectors. Therefore, it is notable that the individual components are temporally and spatially smooth, and the spatial maps are typically left-right symmetric for individual mice. More impressively, the spatial maps are highly similar across different sessions, quantified by measuring the spatial correlation between each pair of sessions for same (‘within component’) and different (‘across components’) components. (Figure 3F). To form a unified map that may highlight which regions are active for each different behavior, we averaged across components that are most active during navigation, timing, or reward (Figure 3G). As expected, visual and somatomotor areas were highly active for navigation and prefrontal areas for reward. For timing, activation was widespread, with a focal region in secondary motor cortex and an unexpected region in barrel cortex. Within entorhinal cortex, the picture is mixed, with all components contributing but with heterogeneity within the area imaged through the microprism.

### Isolating task-relevant and movement-driven signaling

Each task component may be heavily confounded with motor activity, and thus it is possible that signaling during reward consumption, for example, could be due to immobility (mice typically stop running to drink in our task), sensorimotor signaling related to tongue movement and contact with the lick spout, or some intrinsic signal of reward. While it is difficult to separate out completely the contribution of each variable to the observed signals in each pixel, we employed non-negative ridge regression to aid in parsing out these contributions. Predictors included both movement-related predictors and task-related predictors (Figure 4A). Movement-related predictors are calculated from the optical encoder that measures treadmill velocity and from a camera recording the face, which allowed measurement of licking, sniffing, whisking, and pupil diameter. Task-related predictors utilized a basis set based on time relative to the door start and stop (timing basis set) and time relative to reward delivery and teleportation (reward basis set).

**Figure 4.**
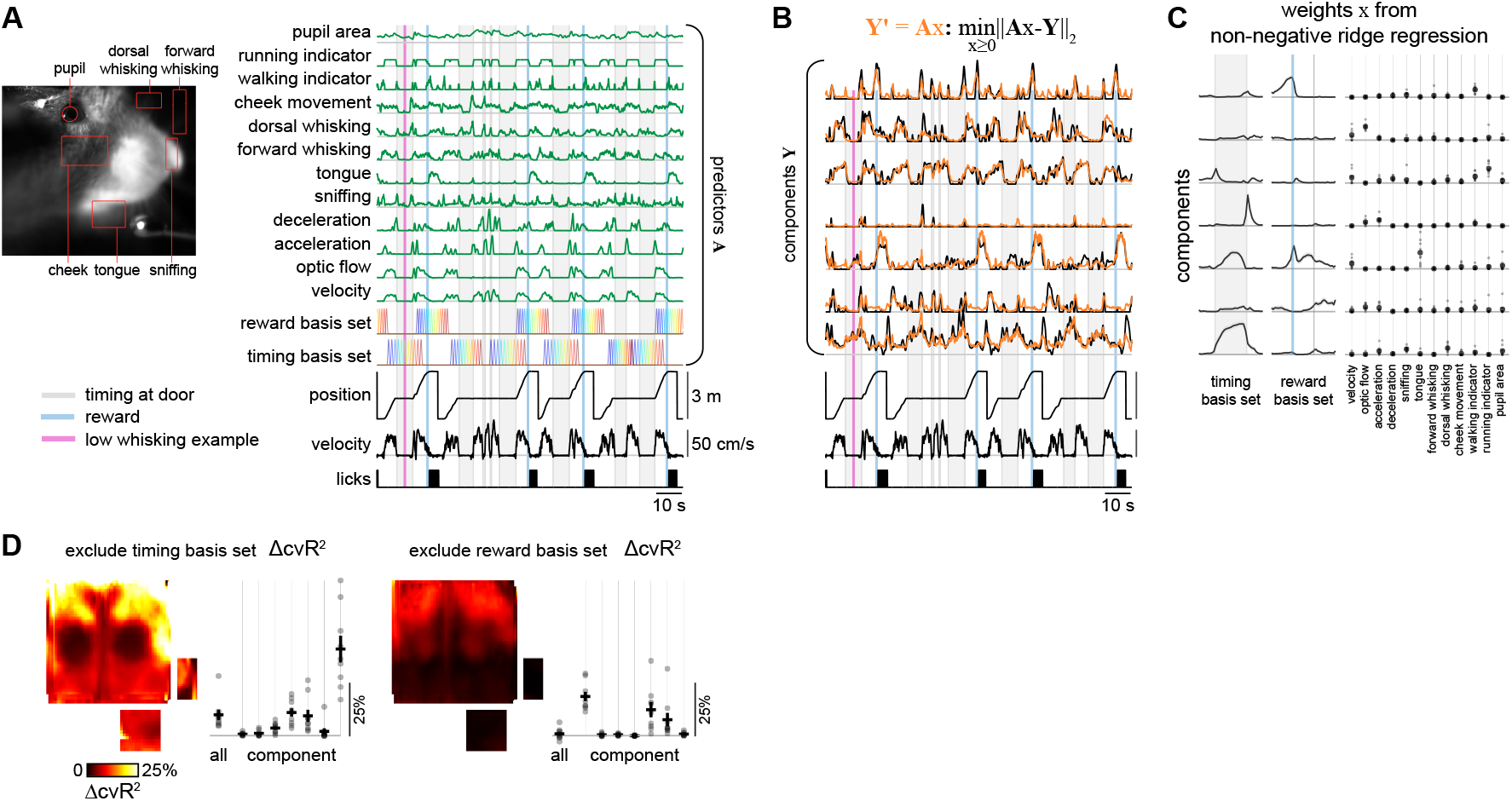
Relation of ongoing behavior to individual components. A) Predictors to be used for non-negative ridge regression, shown for the same exemplar period shown in Figure 2B. These include a timing basis set and a reward basis set, formed by a set of indicator functions at different temporal shifts, here represented by different colors. Behavior predictors are computed from the treadmill velocity (velocity, optic flow, acceleration, deceleration, walking indicator, and running indicator) and from the movie of the mouse’s face (sniffing, tongue movement, forward whisking, dorsal whisking, cheek movement, and pupil area). B) Exemplar traces, projected onto the seven components from Figure 3, along with fit by non-negative ridge regression. C) Weights from non-negative ridge regression for the predictors shown in A, averaged across nine sessions. D) Decrease in cross-validated R^2^ (ΔcvR^2^) when timing basis set (left) or reward basis set (right) is excluded from the predictors used for the regression. Maps show ΔcvR^2^ for each pixel, averaged across *n* = 9 sessions and registered to a common coordinate system. ΔcvR^2^ is also computed for each of the seven components from g-NNMF (light gray dots represent each session, dark cross represents mean ± s.e.m.).

We first tested whether these predictors (**x**) sufficiently capture the signals observed in each region. Time-series vectors for each of the seven components identified by g-NNMF (**Y**) were fit using non-negative ridge regression with 10-fold cross-validation (Figure 4B). The reconstructed signals **Y’** using this approach provided an efficient method of approximating the original movie using a low-dimensional projection, with the reconstructed estimate of the full movie **F’** (formed by multiplying **Y’** by the weights **W**) explaining 79% ± 6% (*n* = 9 sessions) of the variance in **F**. This approach captures similar trends as seen in the temporal vectors for the components as shown in Figure 3D for timing and reward but also serves both to separate contributions from co-mingled variables (such as deceleration and reward approach) and to parcel out contributions from different dimensions of motor activity (Figure 4C). Some of the resulting weights are as expected. For example, component 2, which is prominent in visual cortex (Figure 3E), has its largest weight in the regression for optic flow (Figure 4C). Component 3, which has strong weights in somatomotor cortices (Figure 3E) and closely tracks treadmill velocity (Figure 4B), reassuringly has a strong weight in the regression for running (Figure 4C).

Component 7 is particularly interesting. It would be reasonable to predict that its activation during timing, as reflected in its temporal vector (Figure 3D), would be explained by another variable, such as ongoing whisking or changes in arousal^24^, especially since weights are highest in regions corresponding to barrel cortex. Alternatively, its activation may instead be related to some internal timing process. We located a timing period during which whisking was minimal (Figure 4A, indicated by the vertical magenta line). Interestingly, at this same moment component 7 was highly active, indicating a dissociation from whisking (Figure 4B). This observation is supported by the results of the linear regression. Even though whisking (dorsal and forward) and arousal (via pupil diameter) are predictors, their weights in the regression are not particularly prominent (Figure 4C). Instead, regression weights for the timing basis set itself are prominent. Thus, we conclude that activation of barrel cortex during timing periods is not adequately explained by any externally measurable motor activity and instead is consistent with internal timing.

As the components identified by g-NNMF may be inadvertently discarding some information, we applied the non-negative regression to the raw signals for each individual pixel. The resulting cross-validated R^2^ captured 34.7 ± 3.3% (mean ± s.e.m., *n* = 9 sessions) of the variance^15^. Note that this prediction is being applied to the deconvolved (albeit smoothed) fluorescence trace acquired at 29.4 Hz, so considerable noise is expected. To isolate the contribution of task-relevant variables, we formed reduced models, where we removed either the timing basis set predictors or the reward basis set predictors and measured the resulting decrease in cross-validated R^2^ (ΔcvR^2^, Figure 4D). We also computed the same variable for the individual g-NNMF components. Two results are readily apparent. First, the timing basis set contributes much more to the model than the reward basis set, although this result is partially obscured by the inclusion of tongue movement as a predictor, which highly correlates with reward consumption. Second, the spatial localization of the error map when excluding the timing basis set shows hot spots in barrel cortex and areas of prefrontal and premotor cortex, which provides a complementary approach to g-NNMF for localizing timing-related regions in cortex (and confirmed by the large ΔcvR^2^ for component 7).

### Timing subnetworks dominate as expertise in timing is learned

Given the known task-dependence of neocortex^25,26^, we wondered if task-related changes in cortex occur during learning of the timing task. We divided our sessions into “novice” and “expert” cohorts based on their wait time distributions following either an exponential or Gaussian distribution, respectively (Figure 5A). We verified that there were no obvious differences in movement between these two groups (Supplementary Figure 4). Removing the timing basis set from the linear regression led to a large decrease in variance explained (Figure 5B), indicating that cortical signals in expert mice are driven by timing itself and not correlates of movement. To explore further the cortical signals during timing, we examined the regression weights for timing (timing basis set) in novice and expert mice (Figure 5C). Two features are readily apparent. First, weights are larger in expert mice across the entirety of the timing interval. Second, regions with large weights are distributed across entorhinal cortex, barrel cortex, prefrontal cortex, and retrosplenial cortex, and each peaks at a different point within the timing interval.

**Figure 5.**
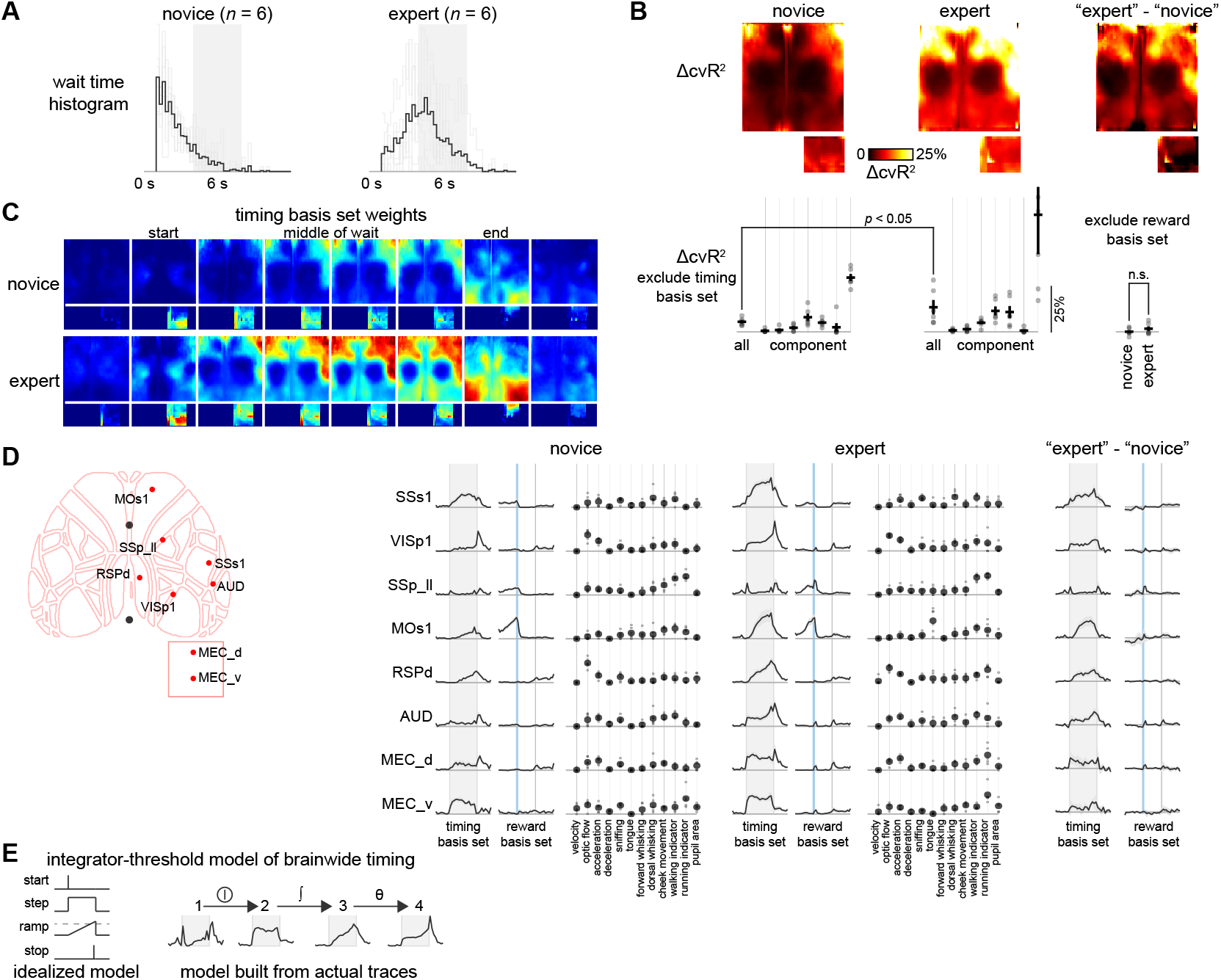
Learning enhances signaling of timing period in cortex. A) Imaging sessions are divided into novice and expert sessions based on the histogram of wait times following an exponential or Gaussian distribution, respectively. Individual session histograms shown in light gray; mean across six sessions shown in black. Target wait times varied from 4 to 8 s as indicated by gray shaded rectangle. Histograms use 0.25-s bins with short waits (less than 1 s) excluded. B) Decrease in cross-validated R^2^ (ΔcvR^2^) when timing basis set is excluded from the predictors used for the regression. Maps show ΔcvR^2^ for each pixel, averaged across *n* = 6 sessions and registered to a common coordinate system. ΔcvR^2^ is also computed for each of the seven components from g-NNMF (light gray dots represent each session, dark cross represents mean ± s.e.m.). Difference map (“expert” – “novice”) shown on right. Average ΔcvR^2^ also shown for excluding reward basis set. C) Regression weights for the timing basis set predictors shown at eight different lags relative to start and end of timing period for novice and expert sessions. Weights from non-negative ridge regression shown for candidate regions for either novice or expert sessions (n = 6 for each). D) Weights from non-negative ridge regression for indicated pixels, averaged across six novice or six expert sessions. E) Proposed timing model. Idealized model (left) shows proposed computation involving integration followed by thresholding. Similar model is built from actual traces (right). Region 1 is an on/off switch that turns on a step in region 2 during a timing period, which in turn is integrated by region 3, resulting in a ramp. Region 4 then takes a nonlinear threshold of this ramp to produce a spike at the target waiting duration.

To investigate these trends in more detail, we examined regression weights for pixels chosen from various exemplar cortical regions during timing and reward. As expected from the comprehensive maps, individual regions show different time courses along with learning-dependent increases, with ramping in retrosplenial and secondary motor cortices, step functions in barrel and entorhinal cortex, and mixed patterns in visual and auditory cortex (Figure 5D). These findings are consistent with recruitment of multiple regions during learning, and the diversity of signals may support different components of a timer circuit^27^. We propose a simple model that could perform interval timing from a combination of these signals by integrating a step function to produce a ramp that is then passed through a threshold function (Figure 5E), similar to previously proposed models for interval timing in neural circuits that employ integration^28^.

## Discussion

A challenge in ascertaining the functional specialization of any brain region is choosing stimuli and behaviors that optimally engage that brain region. Further, representations of these variables will depend heavily on the task the animal is engaged in^25^. While the behaviors tested here are certainly not comprehensive, the incorporation of a timing task along with spatial navigation and reward helps to engage and isolate multiple dimensions of behavior. Using this approach, we find that distinct regions of neocortex and entorhinal cortex are highly engaged across timing, reward, and navigation. These form potential subnetworks, which are distributed but dedicated for certain dimensions of a task. This finding is in agreement with other studies that have found that many regions of neocortex are not just multi-modal, they are also multi-purpose. For example, neurons in visual^29^, somatosensory^30^, and other cortices^31^ can represent spatial position, and neurons in various regions of cortex can represent time^13,16,32^. Whether and how neurons across regions may work together to track time, reward expectation, or spatial position is not answered by the present study, but our results are consistent with the idea that multiple regions of cortex work together to generate and maintain representations of behaviorally relevant variables.

These findings are enabled by our development of a multifocus mesoscale microscope. This imaging system overcomes one of the common limitations of conventional microscopy: a single focal plane. Due to the abundance of photons when imaging modern GFP-based Ca^2+^ sensors such as GCaMP6s in transgenic mice^20^, emission can be split to two separate sensors while still maintaining sufficient signal intensity at both. This approach is used here to achieve separate focal planes^33^ but could also be used for other purposes, such as different emission spectra by employing separate emission filters for each path. Furthermore, we deployed non-negative analysis approaches throughout this study, reflecting the non-negative nature of typical Ca^2+^ signals transmitted by GCaMP sensors, which also benefits from the inherent sparsity that results from non-negative methods^23^. While negative or inhibitory signals are possible with Ca^2+^ imaging^34^, g-NNMF performed similarly to PCA with our fluorescence signals, indicating that the non-negativity assumption is justified. This assumption in turn aided interpretability and separability. For example, using standard linear regression, periods of stationarity could be fit with a negative weight for the running velocity predictor; however, with non-negative regression, weight is put towards the timing basis set.

The identification of a timing subnetwork, which is broadly distributed across much of neocortex and entorhinal cortex, was facilitated by using a behavioral task that requires an interval to be timed explicitly. Further, this timing period requires the mouse to be stationary, which helps isolate timing-related signals from motor-related signals^15^. Surprisingly, the components identified by non-negative factorization revealed that the barrel cortex was a major contributor during timing. This relation was not explained by an increase in whisking during the timing period. One fascinating possibility is that the intrinsic timing circuits that drive rhythmic whisking^35^ could be repurposed to help measure time intervals in trained mice, consistent with previous work showing neurons in somatosensory cortex code for non-sensory task variables – including timing – with learning^36^. Future work will be needed to see if this speculation is indeed true, but this hypothesis aligns with theories that primary sensory neocortex is highly adaptable^26^. Our results are consistent with functional specializations for timing, spatial navigation, and reward across the neocortex that are adaptable to task demands, and these representations may then be communicated via the entorhinal cortex to the hippocampus and other brain regions to inform memory and behavior.

## Methods

All animal procedures were approved by the Northwestern University Institutional Animal Care and Use Committee. Mice expressing GCaMP6s were generated by crossing tetO-GCaMP6s mice (JAX No. 024742) with Camk2a-tTA mice (JAX No. 007004)^20^.

### Surgery

Microprism implants for medial and lateral entorhinal cortex were performed as described previously^21^. Window for transcranial imaging of dorsal neocortex was formed by applying multiple layers of superglue, such as Norland 81, and a large coverslip (either 8 mm square or 10 mm round) placed above. A custom titanium headplate was then affixed using dental cement. Mice were allowed up to a week to recover before starting on water restriction (1 mL per day).

### Behavior

Mice were first habituated to head fixation and running on a linear treadmill. Water rewards were delivered at the end of a 3.1-m virtual track. Daily sessions lasted 40-60 minutes. Once mice ran at least 2 laps per minute, the door stop was introduced, as previously described^13^. Mice were required to reach the door and stop running for a given target interval (initially 1.5 or 2 s) before the door would open, allowing them to continue to the end of the track to receive the water reward. If mice started running before the target interval, the door remained closed and the timer was reset until the next stop attempt. Once mice achieved at least 1 lap per minute, the target interval was increased. For initial training, the door was visible, thus providing a visual cue once the target interval was reached. However, eventually the door was made invisible, requiring the mice to rely completely on an internal clock. Fully trained mice would achieve 1 lap per minute for a 6-s target interval with the invisible door. One to two months of training was typically required before mice reached this level of performance.

### Multifocus Microscope, Imaging, and Hardware

A 50:50 beamsplitter on the emission path split the image to a pair of tube lenses. Distance from each tube lens to each camera was independently controlled. For small deviations, the relationship between the back focal length L (tube lens to camera) and working distance D (objective to imaging plane) can be approximated as L*D = C, where C is a constant that depends on the optics being used. For our purposes, C = 3300 mm^2^. For small deviations near the optimal focal length L, deviations in D are approximately the same but in the opposite direction. Thus, for L = 60 mm, D = 55 mm. For L = 57.4 mm, D = 57.5 mm. Adjusting the back focal length also introduces changes in the magnification, so that decreasing L will increase D and decrease the magnification^33^. But this change in magnification is minimal for small changes in the back focal length. Using this strategy, each camera could be independently focused to a different working distance.

Camera 1 (Photometrics Prime BSI Express) was used as the master, running with an exposure time of 17 msec. Excitation was provided by a pair of LEDs, centered at 470 nm and 405 nm, which were alternatively turned on by a custom-built flip-flop circuit driven by the frame sync from camera 1. Camera 2 was triggered by the frame sync from camera 1, so both cameras ran at 60 Hz while the LEDs pulsed at 30 Hz. Camera 3 was triggered by the same signal that drove the 470 nm LED, so it ran at 30 Hz. All camera and LED times were sent as TTL pulses to an NI-DAQ along with other behavioral data (treadmill velocity, lick sensor data, reward pulses), allowing synchronization of all relevant signals. The collection of data and virtual reality environment were programmed in MATLAB using ViRMEn^37^ and custom scripts.

### Image Processing

For rigid registration across sessions and mice, bregma and lambda were manually chosen from a fluorescence image for each session. A vasculature mask was generated by thresholding for low intensity pixels, and these pixels were subsequently ignored. Images were then downsampled to 0.1 x 0.1 mm^2^ superpixels. Fluorescence traces for 470 nm and 405 nm illumination were formed for each superpixel, and a corrected signal F_C_(t) was calculated by subtracting the dark current and then dividing the 470 nm signal by the 405 nm signal. The baseline F_0_(t) was approximated by estimating weights of a basis set of slowly varying exponentials and sinusoids, allowing calculation of ΔF/F_0_. Next, spatial maps were deblurred using a Lucy-Richardson deconvolution with a σ = 500 μm point spread function^38,39^. Finally, signals were deconvolved^40^ using a double exponential (80 msec rise tau, 500 msec decay tau) to recover an estimate of firing rates.

For the behavior camera (Camera 3), regions of interest were manually selected by drawing rectangles around desired regions (sniffing, tongue movement, forward whisking, dorsal whisking, cheek movement). Energy for each region as calculated as the mean of the square of the temporal derivative (taken for each pixel). For regression analysis, a threshold was applied to remove large spikes, and then this signal was passed through an inverse tangent function. For pupil diameter, we adapted a previously published method that thresholds the pupil region based on intensity and then fits an ellipse to this region^41,42^.

### Analysis

Non-negative matrix factorization was performed using the function nnmf in MATLAB. Non-negative ridge regression was performed using a custom-written script in MATLAB. Quadratic programming with quadprog was used to speed computation of the regression.

---

**Supplementary Figure 1.**
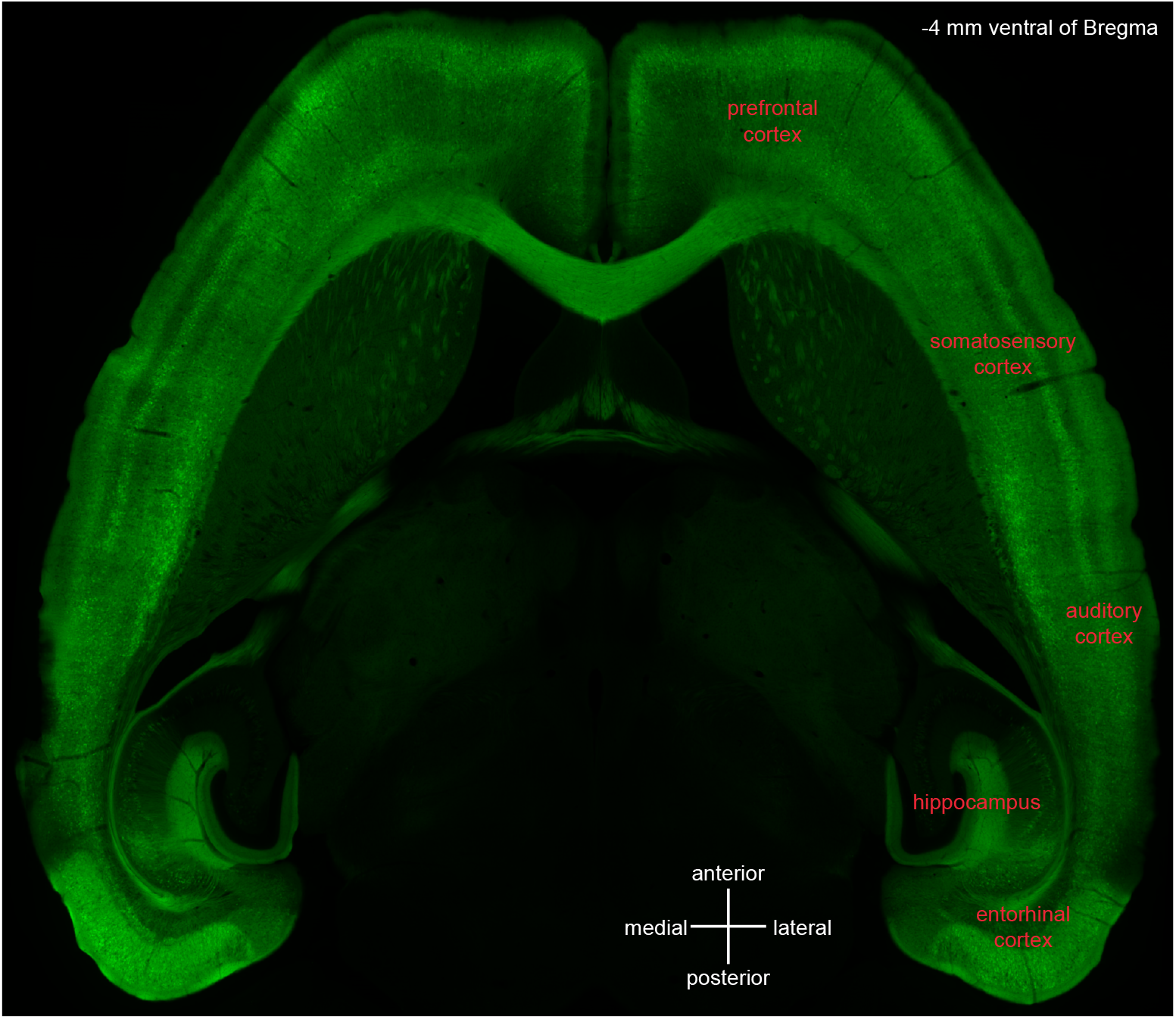
Horizontal slice of transgenic GCaMP6s mouse. Transgenic GCaMP6s mouse was perfused and 50 µm horizontal slices obtained. Image of slice obtained 4 mm ventral of Bregma showing widespread GCaMP6s expression throughout the neocortex and the entorhinal cortex.

**Supplementary Figure 2.**
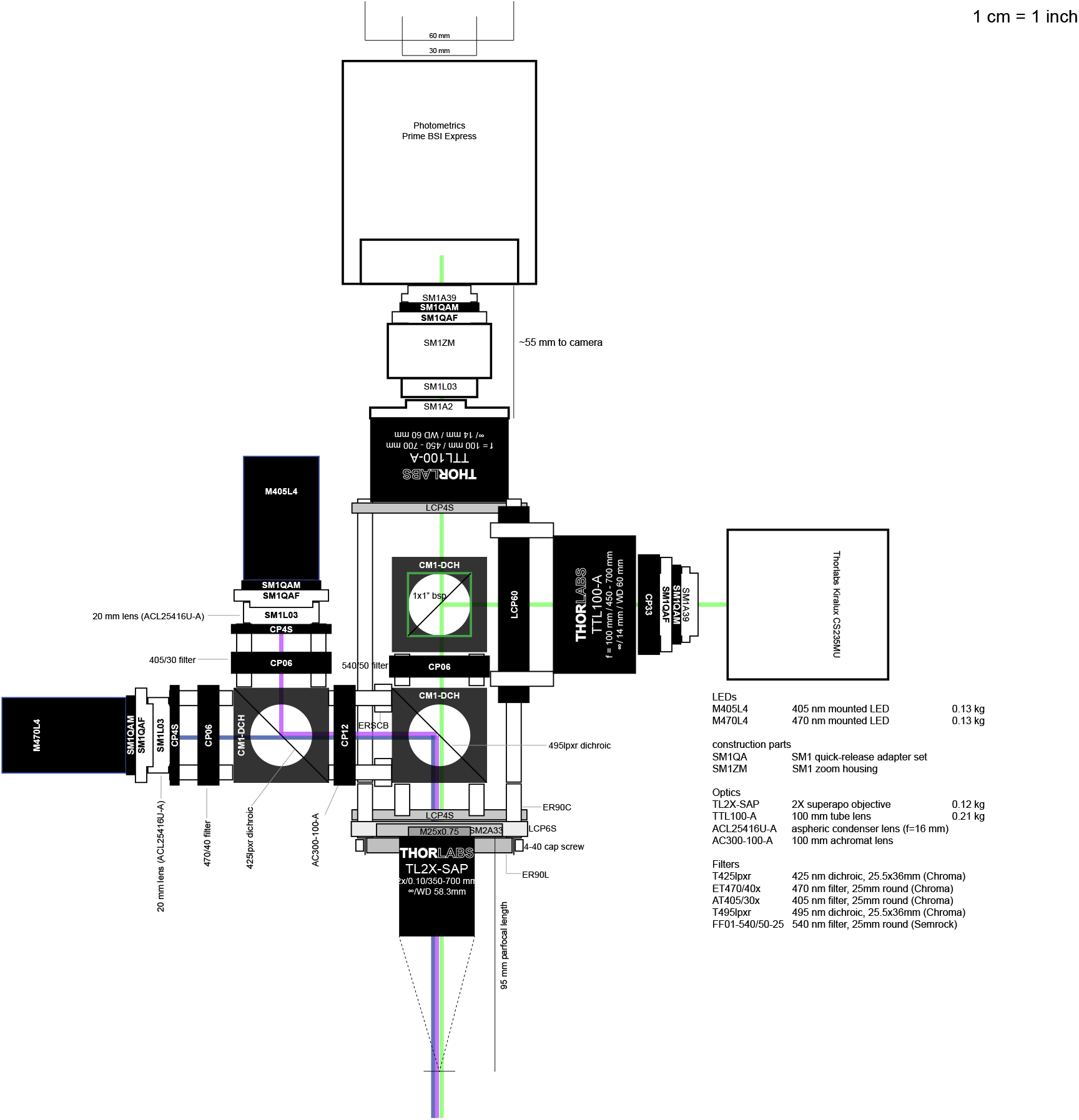
Detailed diagram of multifocus microscope. Detailed view of multifocus microscope. Parts are from Thorlabs unless otherwise indicated.

**Supplementary Figure 3.**
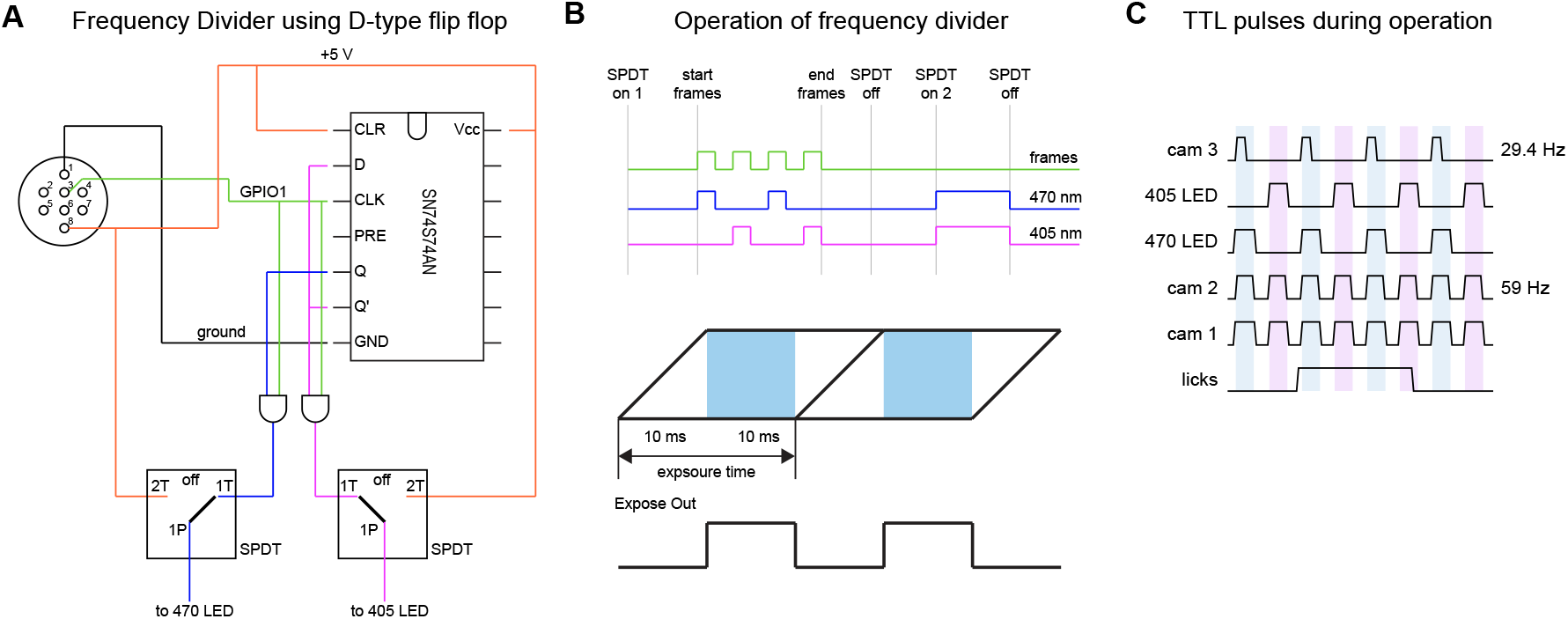
Circuit used to alternate illumination sources. A) Design of flip-flop circuit to perform frequency division. B) Operation of circuit showing alternating TTL pulses delivered to 470 nm and 405 nm outputs. Because the cameras utilize rolling shutters, we avoid exposure across overlapping frames by only exposing when all rows for a frame are being exposed (“Expose Out”). C) Actual operation for the microscope, with three cameras and two light sources. Camera 1 (used as the master) produces alternating TTL pulses used to toggle between the two light sources (405 nm LED and 470 nm LED) and also drives camera 2 (full frame rate) and camera 3 (half frame rate).

**Supplementary Figure 4.**
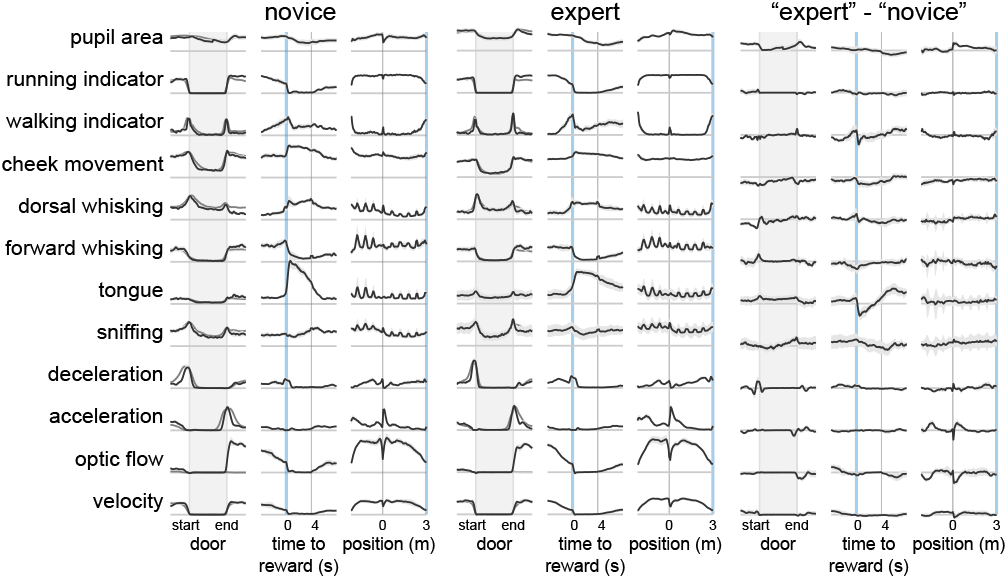
Change in motor activity as expertise is gained in the timing task. Triggered averages of different read-outs of motor activity in novice and expert mice.

